# Quantitative comparison of the anterior-posterior patterning system in the embryos of five *Drosophila* species

**DOI:** 10.1101/378430

**Authors:** Zeba Wunderlich, Charless C. Fowlkes, Kelly B. Eckenrode, Meghan D. J. Bragdon, Arash Abiri, Angela H. DePace

## Abstract

Complex spatiotemporal gene expression patterns direct the development of the fertilized egg into an adult animal. Comparisons across species show that, in spite of changes in the underlying regulatory DNA sequence, developmental programs can be maintained across millions of years of evolution. Reciprocally, changes in gene expression can be used to generate morphological novelty. Distinguishing between changes in regulatory DNA that lead to changes in gene expression and those that do not is therefore a central goal of evolutionary developmental biology. Quantitative, spatially-resolved measurements of developmental gene expression patterns play a crucial role in this goal, enabling the detection of subtle phenotypic differences between species and the development of computations models that link the sequence of regulatory DNA to expression patterns. Here we report the generation of two atlases of cellular resolution gene expression measurements for the primary anterior-posterior patterning genes in *Drosophila simulans* and *Drosophila virilis*. By combining these data sets with existing atlases for three other *Drosophila* species, we detect subtle differences in the gene expression patterns and dynamics driving the highly conserved axis patterning system and delineate inter-species differences in the embryonic morphology. These data sets will be a resource for future modeling studies of the evolution of developmental gene regulatory networks.

## Introduction

In the embryo, naïve cells are patterned into complex tissues by precise programs of gene expression that unfold over developmental time. A cell’s eventual fate is determined by the spatiotemporal expression patterns of key patterning genes. Therefore, a change in embryonic gene expression patterns can drive divergence of an organism’s adult form, and conversely, conservation of gene expression patterns, despite changes in the regulatory DNA that encodes them, can maintain a developmental program over large evolutionary distances (Carroll et al., 2005; Davidson, 2006; Gordon and Ruvinsky, 2012; Halfon, 2017; Rebeiz and Tsiantis, 2017).

Early embryogenesis in Drosophila provides an interesting case study for the evolution of development. Its axis patterning systems are qualitatively conserved across the genus, in spite of 40 million years of evolution and sequence diversity in the coding regions equivalent to that of all amniotes (Lin et al., 2008). This patterning system is deployed in Drosophila species that develop under differing conditions of temperature, humidity, and atmospheric composition, which may affect the embryo’s physical characteristics and constraints (Ashburner et al., 2011).

To understand how regulatory DNA sequences encode developmental programs and to detect subtle evolutionary differences in embryonic patterning, there is a need for quantitative, spatially resolved measurements of the expression patterns of developmental genes across species. For example, comparisons of early axis patterning between *Drosophila melanogaster* and the scuttle fly yielded insights into how a common developmental program was conserved, despite system drift (Wotton et al., 2015). Extensive work comparing the regulatory network that defines the endomesoderm in several species of sea urchins has revealed network motifs that meet patterning challenges (Hinman and Cheatle Jarvela, 2014). Ideally, gene expression measurements would be made at cellular resolution (since this is the natural unit of measure in the organism), for all the relevant genes in a patterning system, using a uniform technique across species, and reported in an easily shared format.

Here we report the generation of two gene expression atlases for *Drosophila simulans* and *Drosophila virilis*, which include cellular-resolution measurements of the core anterior-posterior patterning genes in the early embryo. By combining these data sets with existing measurements for *D. melanogaster* (Fowlkes et al., 2008), *D. yakuba*, and *D. pseudoobscura* (Fowlkes et al., 2011), we compared gene expression patterns across five species of *Drosophilids*, spanning 40 million years of evolution. We identified differences in the gene expression patterns between species and the embryo sizes, shapes, and nuclear numbers and provide these data sets in an easily distributed format for future modeling studies.

## Results

*Generation of gene expression atlases for two species in the early embryo* We generated cellular-resolution gene expression atlases for *D. simulans* and *D. virilis* spanning six time points during the blastoderm stage of embryogenesis. These atlases were made using the same methodology as existing *D. melanogaster*, *D. yakuba* and *D. pseudoobscura* atlases (Fowlkes et al., 2008; Fowlkes et al., 2011) and provide cellular-resolution measurements of average gene expression for 13 core anterior-posterior (AP) patterning genes in *D. simulans* and 10 AP genes in *D. virilis. Caudal (cad), forkhead (fkh)*, and *paired (prd)* were not included in the *D. virilis* atlas because we were unable to generate probes that yielded quality *in situ* hybridization patterns. Due to the differences in the mRNA and protein patterns, we also measured hunchback (hb) protein levels in *D. simulans, D. pseudoobscura, D. yakuba*, and *D. virilis* to complement the existing *D. melanogaster* data (Figure 1). To generate each atlas, embryos were stained for a gene of interest, a fiduciary marker gene, and DNA, and then imaged and processed into “pointclouds,” text files that contain the spatial coordinates for each nucleus and the level of expression for each stained gene. Using the fiduciary marker, each embryo was aligned to a species-specific morphological template, which allows several embryos stained for the same gene to be averaged, and stains for multiple genes of interest to be combined (see Table S1, Methods).

**Figure 1.**
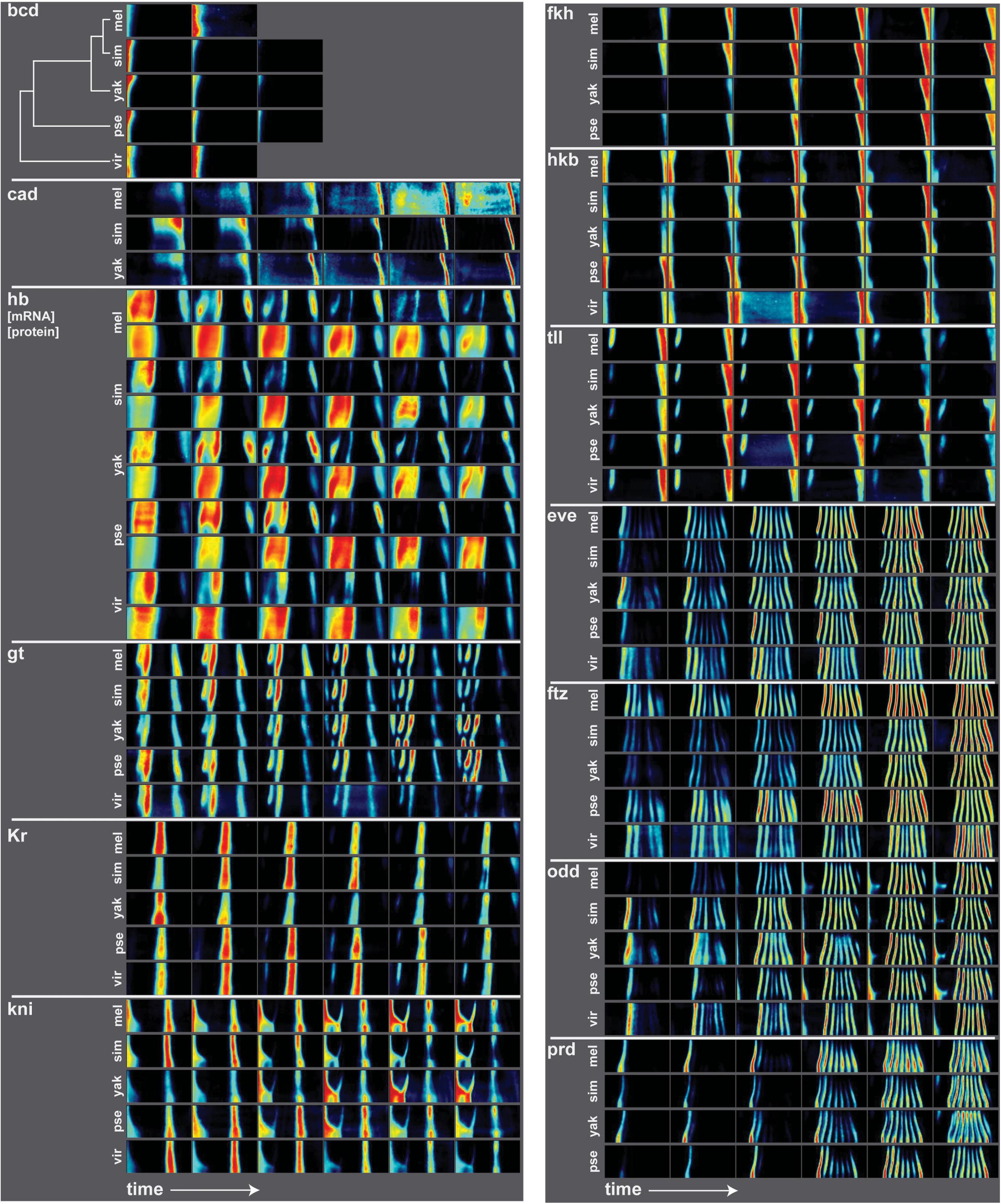
Average gene expression patterns for segmentation genes in five species. Here we show the average gene expression patterns for key maternal, gap, terminal and pair rule genes from the *D. melanogaster* (Fowlkes et al., 2008), *D. yakuba, D. pseudoobscura* (Fowlkes et al., 2011), *D. simulans*, and *D. virilis* atlases. Gene expression is depicted as a heat map, with black corresponding to no expression and red corresponding to high expression. The patterns are shown as “unrolled” half embryos, since the patterns are left/right symmetric. In this depiction, anterior is left, posterior is right, dorsal is up and ventral is down, and developmental time is increasing from left to right. The patterns are qualitatively similar, but vary quantitatively in their dynamics and the relative intensity of different parts of the expression patterns.

As with previous atlases, we defined six time points within the blastoderm stage using a morphological marker instead of clock time. The developmental timing of these species varies considerably (Kuntz and Eisen, 2014), so comparable developmental stages are more easily identified using a morphological marker. During this stage of development, the syncytial embryo becomes cellularized, so we used percent membrane invagination to determine developmental stage and divided the embryos into time points that correspond to roughly 10-minute intervals in *D. melanogaster* (Fowlkes et al., 2008) (see Methods).

The *D. simulans* and *D. virilis* atlases are of similar qualities to the previously measured atlases. To assess the quality of these atlases, we calculated the average standard deviation of expression values between the embryos used to generate the atlases for each gene and time point and present the values, averaged across time points, in Table S2. Compared the *D. melanogaster* values, 12 of 13 genes in the *D. simulans* atlas and 6 of the 10 genes in the *D. virilis* have lower standard deviations.

### Qualitative differences in gene expression patterns

As expected, the patterns of expression of these highly conserved patterning genes are qualitatively similar between the species, but there are several subtle differences. In the gap genes, there are several species-specific patterns, especially in the anterior expression domains. For example, In *D. virilis*, the anterior pattern of *giant (gt)* expression differs from the other species in the last two time points, where shows a weaker anterior-most domain of expression. It is possible that this change in *D. virilis gt* expression has functional consequences, since the lack of *gt* causes defects in head structures in *D. melanogaster* (Mohler et al., 1989). The anterior domain of the hb protein pattern shows different dorso-ventral modulation in different species. In later timepoints, the anterior domain splits into two stripes; the dynamics and relative strengths of these stripes are species-specific. In *D. virilis, Kruppel (Kr)* also shows a different pattern of expression. There is a more distinct region of anterior expression, and the species also lacks the posterior expression domain in late time points. The *knirps* (*kni)* expression pattern in *D. virilis* lacks the partial stripe at around 40% egg length from the anterior.

The terminal and pair-rule genes also vary between species. For example, the *tailless (tll)* pattern fades more quickly in *D. simulans* than other species. And the dynamics and relative strengths of pair-rule genes also vary. For example, the relative strengths and dorso-ventral modulation of the *even skipped* (eve) stripes at the final time point differs between species. *Odd skipped (odd)* stripe 7 is very weak in *D. virilis* compared to the other four species, and *D. melanogaster* has a weaker stripe 1 than the rest of the species.

### Blastoderm embryos vary in nuclear number, shape and nuclear density

To generate gene expression atlases, an average morphological embryo template must be generated for each species, which can also be used to study the morphology of the blastoderm embryo itself (Table 1, Figure 2). Among the five species studied, *D. yakuba* has the highest average nuclear number, followed by *D. melanogaster, D. simulans, D. virilis* and then *D. pseudoobscura. D. virilis* is the longest of the five species we have measured, and *D. yakuba* is nearly as long, but has many more nuclei than *D. virilis*, suggesting that that *D. virilis* cells are larger than the other species.

**Figure 2.**
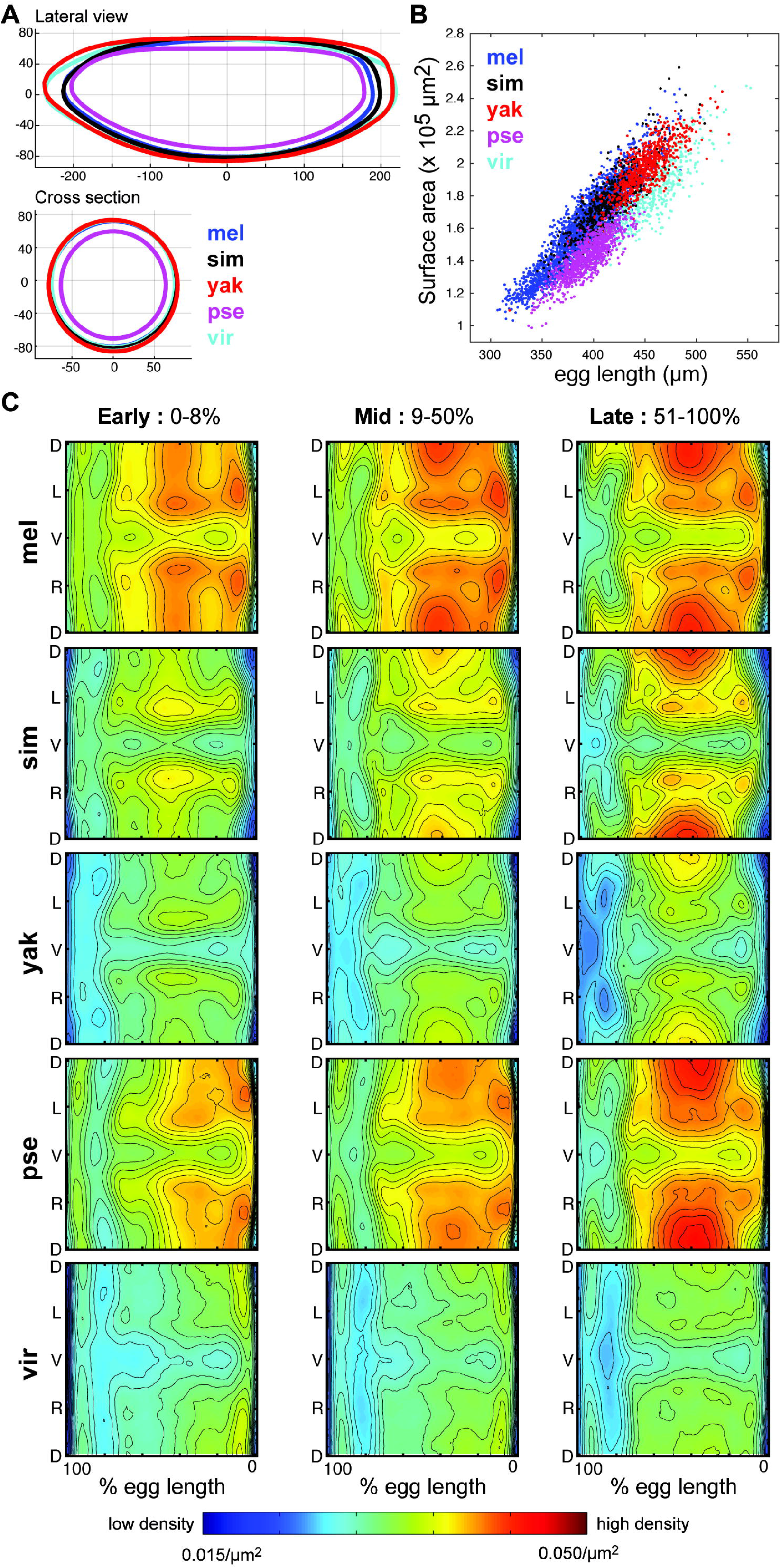
Embryos vary in shape and size. (A) The lateral and cross-sectional views of the average embryo shape show that *D. pseudoobscura* embryos are the smallest, followed by the similarly sized *D. melanogaster* and *D. simulans* embryos, and then the larger *D. virilis* and *D. yakuba* embryos. Units are in microns. (B) All embryos show a similar scaling between surface area and embryo length, indicating that the average shape of the *Drosophilid* egg is conserved, and uniformly stretched or contracted between the species. (C) Here we show nuclear density patterns for all five species at three time intervals in the blastoderm stage of development: early (0-8% cellular membrane invagination, time points 1-2 from Figure 1), mid (9-50% invagination, time points 3-4), and late (51-100% invagination, time points 5-6). All show the characteristic low density patterns in the regions where the cephalic and ventral furrows will form during gastrulation, but average density varies considerably from one species to the next, indicating that embryos with similar surface areas (e.g. *D. yakuba* and *D. virilis)* have different nuclear numbers.

**Table 1.** Average nuclear number and egg length for five species

| **Species** | **Embryos** | **Ave. No.**<br>**Nuclei** | **Std. Dev.** | **Ave. Egg**<br>**Length**<br>**(um)** | **Std. Dev.** |
| --- | --- | --- | --- | --- | --- |
| *D. melanogaster* | 2772 | 5974.1 | 339.12 | 393.8 | 30.75 |
| *D. simulans* | 613 | 5894.3 | 265.42 | 416.2 | 24.97 |
| *D. yakuba* | 672 | 6114.8 | 342.39 | 450.9 | 22.73 |
| *D.*<br>*pseudoobscura* | 966 | 5081.0 | 327.97 | 394.1 | 18.93 |
| *D. virilis* | 476 | 5535.2 | 436.09 | 457.4 | 26.90 |

As with the other species, patterns of nuclear density in *D. simulans* and *D. virilis* prefigure the cell movements that occur during gastrulation (Blankenship and Wieschaus, 2001; Luengo Hendriks et al., 2006), with regions of low density in the regions that will become the cephalic and ventral furrows (Figure 2C). These patterns of nuclear density highlight that *D. virilis* has the least dense nuclear packing of the five species under study, while *D. melanogaster* and *D. pseudoobscura* have the densest packing patterns.

Measurements of the shape of the embryos also reveal differences between species. *D. melanogaster* and *D. simulans* have similar dimensions, as do *D. virilis* and *D. yakuba*, with *D. pseudoobscura* showing a distinctly smaller circumference and lateral outline (Figure 2A). The relationship between the embryo length and surface area of all five species is quite similar and linear (Figure 2B), reflecting that differences in the surface areas of the embryo both within and between species are largely accounted for by differences in the egg length.

Together these observations confirm that general morphological features of the embryo are conserved, e.g. the similar patterns of low and high nuclear density, but specific properties like nuclear number and embryo size vary quantitatively between species.

### Binary cell type analysis reveals quantitative differences in the patterning network dynamics between species

To assess the similarity of and differences between gene expression atlases, we define cell types based on the combination of genes expressed in the nucleus at a particular time point, as this gene expression profile prefigures the cell’s eventual fate (Lehmann and Frohnhöfer, 1989; Lawrence, 1992; St Johnston and Nüsslein-Volhard, 1992). In this analysis, we use a threshold to determine whether a gene is “off” or “on” in a nucleus and define cell types as the combinations of genes that are on each nucleus. Though this threshold-based approach does not account for quantitative changes in mRNA levels, it is not clear that quantitative changes are sufficient to drive cell fate differences in this network, and our group has previously used this approach to explore the canalization of cell fate in *bicoid* (*bcd*)-depleted *D. melanogaster* embryos (Staller et al., 2015a).

We first considered the nine genes that were measured in all five species atlases in all six time points. This set is composed of all four gap genes: *hb, gt, kni*, and *Kr*, two terminal genes: *tll* and *huckebein* (*hkb*), and three primary pair-rule genes: *eve, odd* and *fushi tarazu (ftz)*. After discarding cell type combinations that are observed in less than 0.1% of the total nuclei, we find that of the 2^9^ = 512 possible cell types, only 79 are actually observed and the fraction of observed cell types decreases with increasing gene number (Table 2). Of these 79 observed cell types, there is only one that is not observed in all five species – it is “*gt hb kni*,” which is missing in *D. virilis*, but only accounts for 0.17% of the total nuclei. Given the high level of conservation of the patterning network between species, we would not expect unique cell types in species, and this confirms that our cell type definitions are generally sound.

**Table 2.**
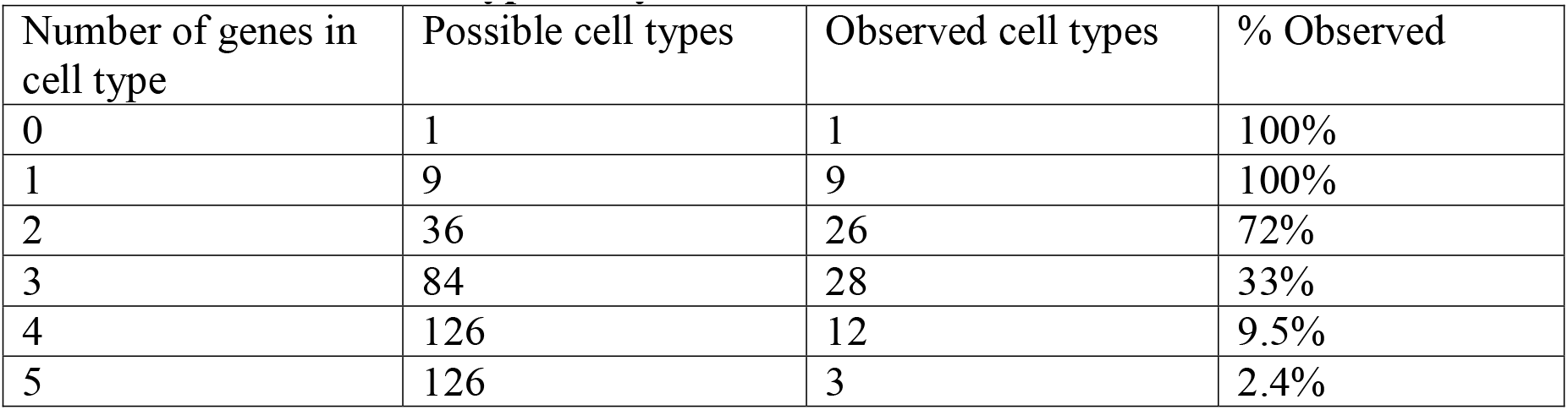
Statistics of cell type analysis

Since our gene set is composed of transcription factors that largely repress one another (Figure 3A), we expect that each nucleus will only express a subset of the 9 genes. The maximum number of genes expressed in a single nucleus is five, but the bulk of nuclei express 1, 2 or 3 genes from our set (Figure 3B). Additionally, the proportion of nuclei expressing 4 or 5 genes decreases markedly over developmental time, which is consistent with the observation that the cross-repression of these genes takes place throughout this stage of development (Jaeger, 2011). While the fraction of nuclei expressing 4 genes remains relatively high in the last time point of the *D. virilis* atlas, this is likely due to higher than average background staining of *gt*. Of 557 nuclei expressing four genes in the last time point of *D. virilis*, nearly half are expressing either *ftz/odd/gt/kni* or *ftz/odd/gt/hb*.

**Figure 3.**
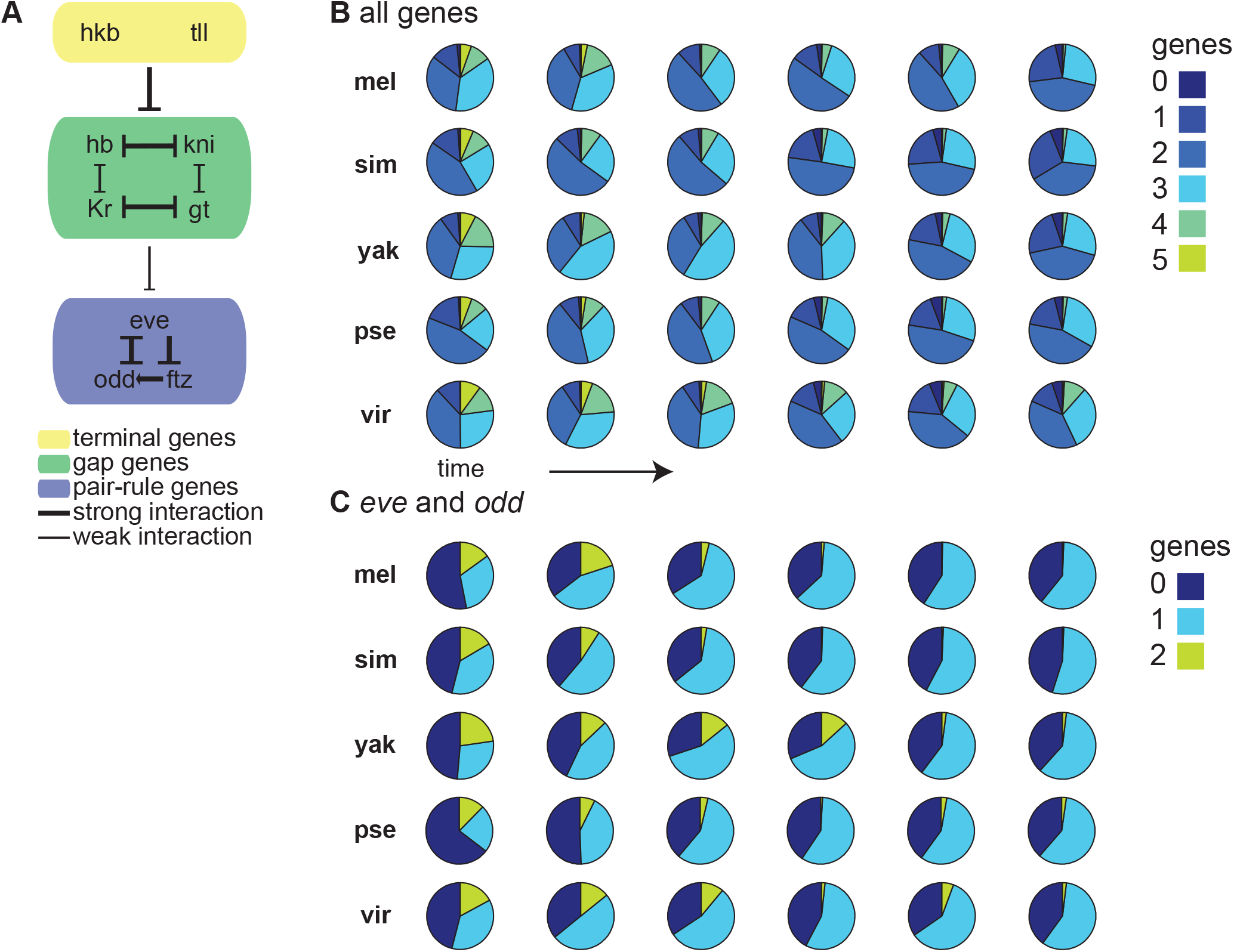
The repressive relationships between genes in the network yield nuclei with fewer genes expressed over time. (A) Among the nine genes measured in all five gene expression atlases, most are repressors. We show the known interactions between these genes with thicker lines indicating stronger interactions and thinner lines indicating weaker or partial interactions, e.g. the gap genes generally repress only part of the pair rule genes’ patterns. (B) These pie charts show the proportion of nuclei expressing 0 to 5 of the genes we assayed as a function of time. When considering the nine genes common to all five atlases, no nuclei expressed more than 5 genes simultaneously. The proportion of nuclei expressing 4 or 5 genes decreases with developmental time, consistent with the idea that the protein products of the repressors accumulate over this developmental time period, reducing the number of nuclei expressing two genes that repress each other. (C) These pie charts show the proportion of nuclei expressing 0, 1 or 2 genes, when only considering *eve* and *odd*. The proportion of nuclei expressing both genes decreases over time, but notably, the number of nuclei with both *eve* and *odd* is quite variable at the first two time points, which reflects differences in the onset of expression for these genes.

To study the dynamics of pair-rule gene expression, we repeated this analysis considering only *eve* and *odd*, and we found that the fraction of cells expressing both *eve* and *odd* decreases over developmental time. This pattern reflects the sharpening of the pair-rule gene expression patterns over time, which is due to the cross-repressive relationships between pair-rule genes expressed in alternating segments of the embryo (Jaynes and Fujioka, 2004; Schroeder et al., 2011). The proportion of cells that express both *eve* and *odd* and the rate at which their cross-repression occurs is species-specific (Figure 3C). This analysis confirms that cross-repression of genes in this system strengthens during the blastoderm stage in all species but shows that the dynamics of the repression differs between species.

### Cell type analysis reveals the drivers of differences between time points and species

We can use the cell type analysis to study the dynamics of cross-repression within a species. In Figure 4A, we cluster the cell type proportions across time points and find distinct clusters of cell types found predominately at each time point in the *D. melanogaster* atlas. Earlier time points have an abundance of nuclei expressing mutually repressive genes, e.g. *eve/ftz* or *hb/kni*, while later time points are dominated by nuclei expressing one gene, e.g. eve, or one gene that is activated by another, e.g. *ftz/odd*.

**Figure 4.**
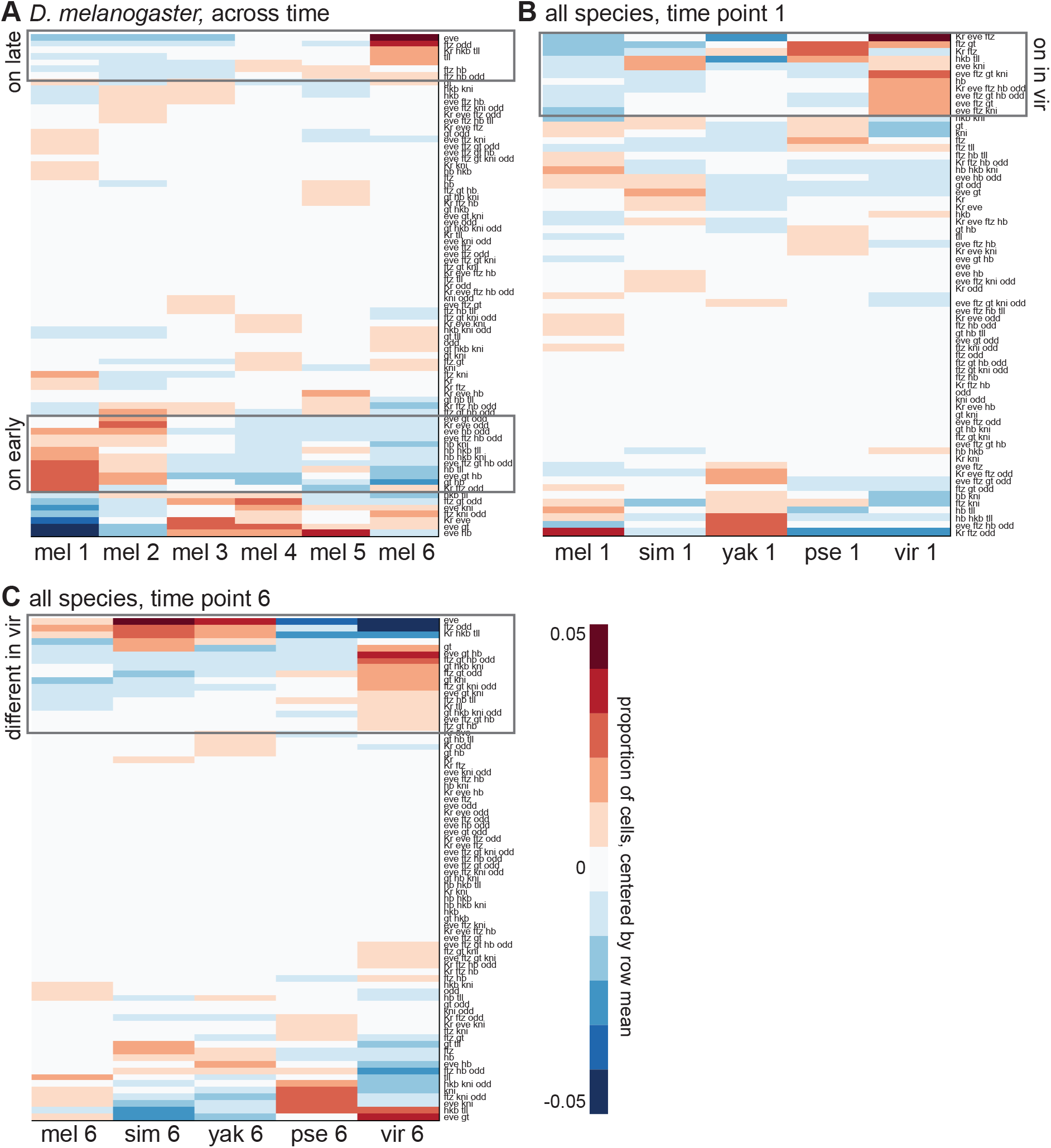
Clustering of cell type profiles reveals the cell types driving differences between time points and species. To explore how the abundance of different cell types change over time or between species, we hierarchically clustered different subsets of the cell type profiles. The rows correspond to cell types and the columns to different species time points. The values shown are the proportion of nuclei expressing a cell type, normalized to the row average, to clearly depict how the proportion is changing over time or between species. The clustergrams show (A) *D. melanogaster*, across time (B) time point 1, across species and (C) time point 6, across species. Gray boxes highlight cell combinations that are differentially present between time points or species.

Lastly, we wanted to identify the changes in the proportions of genes combinations that were most diverged between the species, both at the beginning and end of this developmental time point. In Figures 4B and 4C, we cluster the cell types in time point 1 and 6, respectively. In time point 1, the divergence of *D. virilis* cell types is largely explained by cell types that contain both *eve* and *ftz*, which are due to the relatively strong expression of four fuzzy *eve* stripes in *D. virilis* at time point 1. In time point 6, *D. virilis* has fewer cells with *eve* alone or *ftz/odd* alone than the other species, and a greater proportion of cells with the pair-rules co-expressed with *gt*, which again may be due to the high background of *gt* in time point 6 of *D. virilis*.

### Expression distance scores reflect the underlying phylogeny of the five species

Though the binary cell type analysis is a useful tool for the comparing the general dynamics of the patterning network, it has two limitations: a threshold is needed to define genes as “on” or “off” and reducing each atlas time point into a vector defining the fraction of nuclei of each cell type loses spatial information. To overcome both of these limitations, we employ the “expression distance score,” which was introduced in our analysis of the *D. yakuba* and *D. pseudoobscura* atlases (Fowlkes et al., 2011). Here, each cell is described as a vector containing an entry for its gene expression level for each gene at each time point, and cells are compared by calculating the squared Euclidian distance between these vectors, which we term the expression distance score. We chose to use the squared distance, as compared to Euclidian distance, because it is additive across time points and genes, which makes its interpretation easier.

We first systematically compared cells between the *D. melanogaster* and other species atlases by calculating the expression distance score between each *D. melanogaster* cell and its nearest spatial match in the other species. Since the species’ embryos vary in size, we made spatial matches by normalizing egg length and aligning the embryos’ centers of mass. When these scores are calculated using all nine genes common to the five atlases, two patterns emerge (Figure 5, first column). First, the expression distance score is non-uniform across in the embryo. For example, when comparing *D. melanogaster* to *D. simulans* or to *D. yakuba*, the most divergent cells are found in the anterior section of trunk region. Comparisons to *D. pseudoobscura* or *D. virilis* show more widespread differences throughout the trunk. Second, the average expression distance score between *D. melanogaster* and the other species increases with phylogenetic distance. This increase may be due to an increased divergence in the expression patterns, an increased divergence of the morphological arrangement of cell expression profiles within the embryo, the expansion of contraction of the number of cells with a certain expression profile, or all of the above.

**Figure 5.**
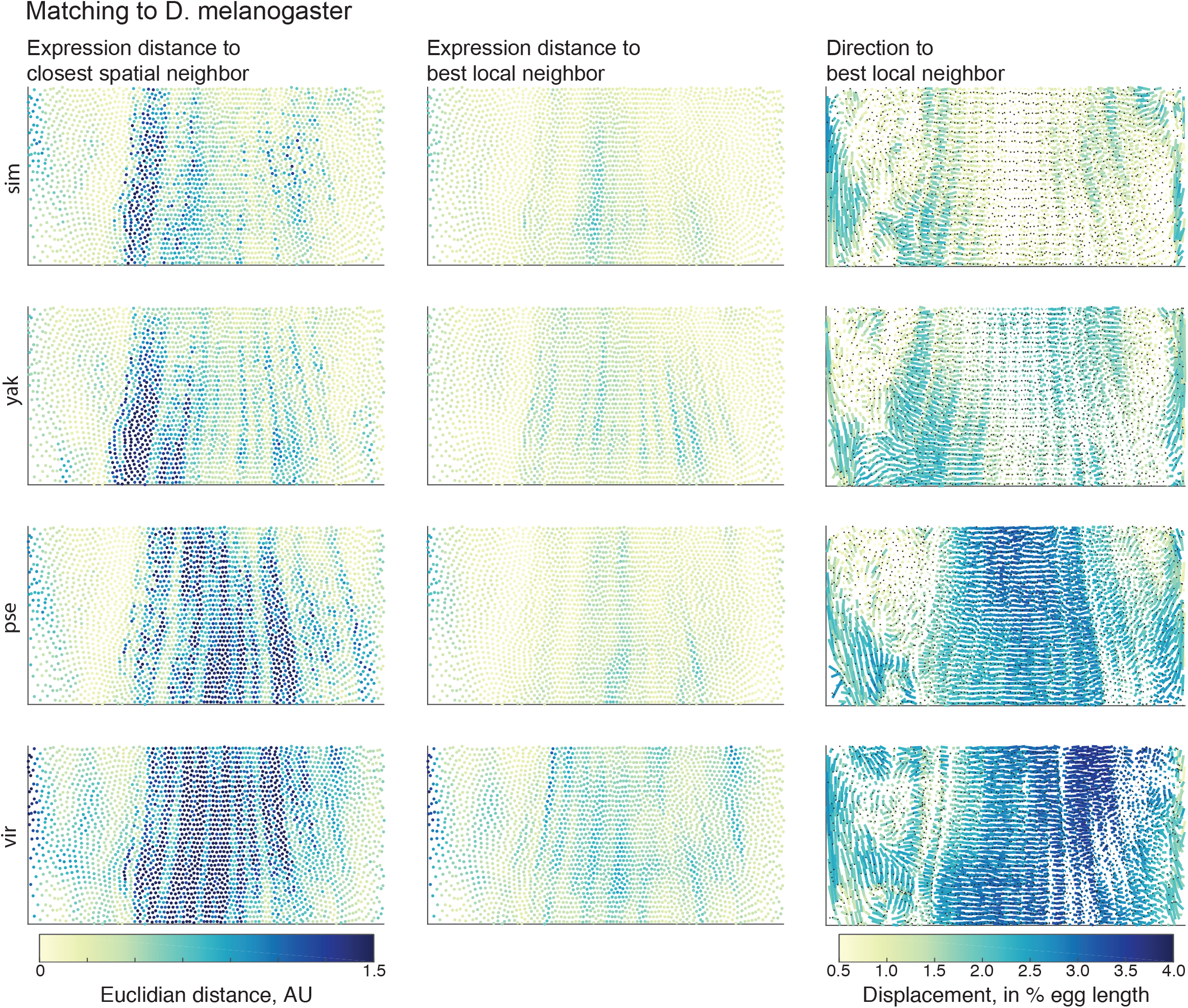
The first and second columns show the expression distance score between *D. melanogaster* and a second species, where cells are either matched to (first column) their nearest spatial neighbor or (second column) the best matching cell within a 30-cell neighborhood. The local search for a best matching cell dramatically reduces the expression distance score, indicating that there generally exist similar cells in the respective atlases, but they may have changed in their exact position in the embryo and/or in their relative abundance. The third column shows the direction and magnitude of distance between matching cells in *D. melanogaster* and another species. The lines connect the location in *D. melanogaster* and the second species, with the black dot indicating the location in the second species. The color of the line indicates the magnitude of the move in 3D space, which may not correspond to the length of the line in this 2D projection.

To remove the effects of morphological and cell proportion differences, we use a second instantiation of the expression distance score. In this iteration, instead of matching each cell to its nearest spatially matching cell in the second species, we conduct a local search, allowing a query cell to be matched to one of its 30 nearest neighbors with the lowest expression distance score. This local matching allows us to remove the influence of the movement of the best matching cell by 3-4 cells in any direction. Because we do not require one-to-one matching, which is challenging and potentially misleading in species with differing cell numbers, this also accounts for expansions or contractions of the number of cells with a certain expression profile. The local matching dramatically decreases the expression distance scores, with the most dramatic decreases in median distance in the *D. melanogaster-D. pseudoobscura* and *D. melanogaster-D. virilis* comparisons (Figure 5, middle column, Figure S1). Even after this matching algorithm, there are nuclei in the anterior region of *D. virilis* that still show larger expression distances, which reflect the qualitative differences in gap gene expression noted in Figure 1. This result indicates that part of the expression distance score divergence can be attributed to coordinated changes in cell expression profile’s morphological arrangement, rather than the lack of a similar cell in each species.

This analysis also allows us to visualize both how far the best matching cells are from one another and the direction of their movement, relative to each other (Figure 5, last column). When comparing *D. melanogaster* to the closely-related *D. simulans*, the magnitude of movement is relatively small, with movements concentrated at the anterior and posterior ends of the embryo and the anterior edge of the trunk region, where there is generally an anterior shift of *D. simulans* cells relative to *D. melanogaster* cells. *D. yakuba* shows a similar pattern, though the magnitude of the movements is generally greater. In *D. pseudoobscura* and *D. virilis*, cell movements are widespread throughout the embryo’s trunk, and the cell shifts are generally posterior relative to the matching *D. melanogaster* cells. The average magnitudes of these shifts again reflect the underlying phylogenetic distances between species.

### Developmental time measured by morphological and gene expression markers diverge between species

We considered a hypothesis that may explain why the dynamics of expression vary from one species to the other. Since the total time of embryogenesis of these species varies (Kuntz and Eisen, 2014), we matched time points by using a morphological marker of the developmental time – the percentage invagination of the cell membrane as the embryo goes from a syncytium to a cellularized blastoderm. It is possible that the gene expression dynamics and membrane invagination do not progress at same rate in each species and that, based on gene expression alone, we can define a different matching between the time points of different species. To see if this is the case, we described the gene expression pattern of each species at each time point as a vector containing 79 values corresponding to the proportion of nuclei of each cell type, as defined by the “on” or “off” calls of the nine genes common to all atlases. We then calculated the Euclidean distance between each pair of vectors and display the resulting values in Figure 6A. To match time points from *D. melanogaster* to each other species, we traversed a path along the distance matrix, starting by matching time point 1 in *D. melanogaster* to time point 1 in the other species, and for each subsequent time point, finding the time point in the other species that had the minimum distance, while not allowing steps “backwards” in time.

**Figure 6.**
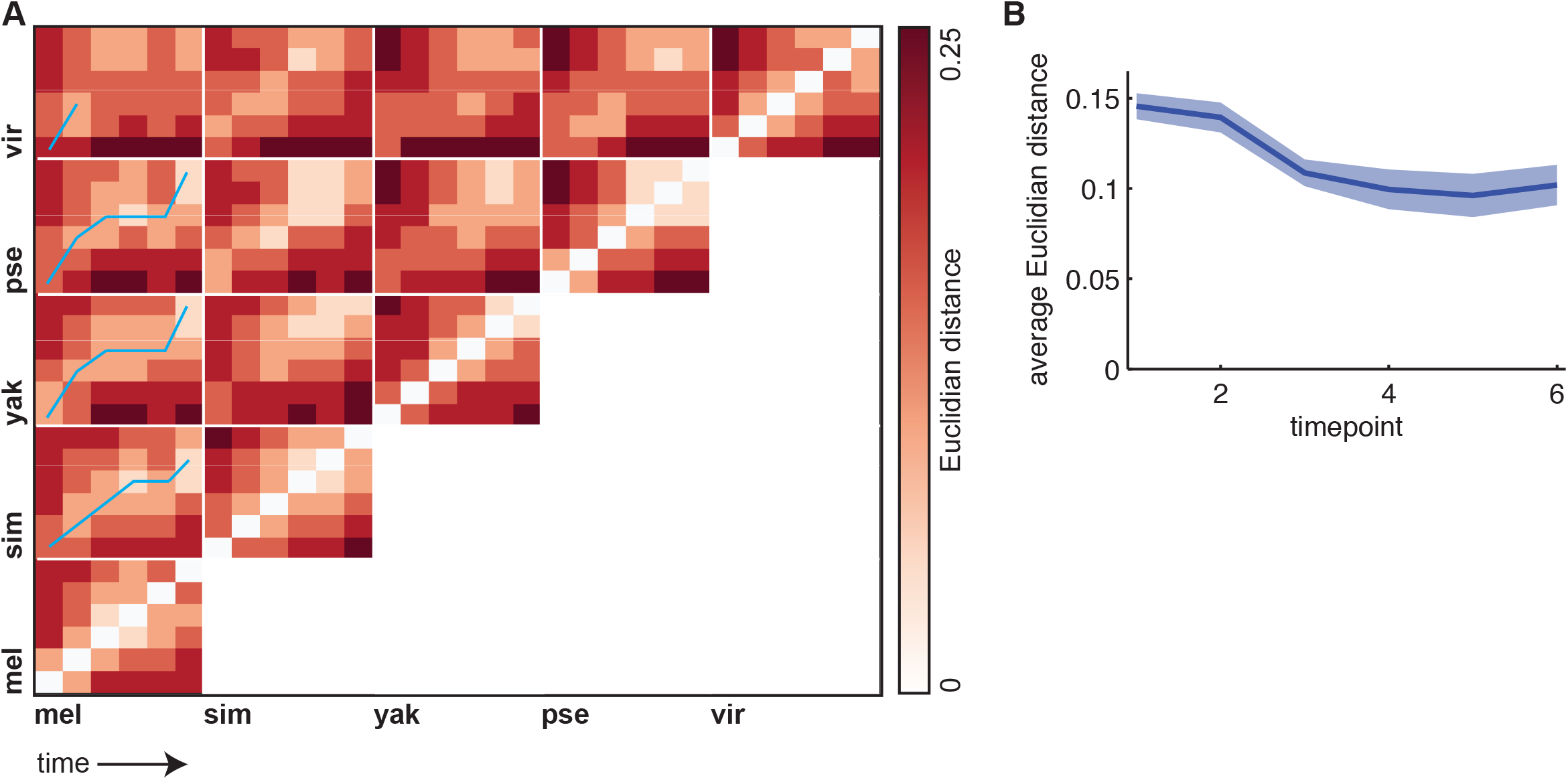
Developmental time as defined by morphology and expression differ between *Drosophilids*. To test whether the dynamics of gene expression patterns match the morphological landmarks used to define the atlas time points, we calculated the Euclidian distance between the proportion of nuclei expressing each combination of genes between each species time point. Smaller distances indicate a greater similarity between the proportions (white). To match time points by cell type profiles, we *matched D. melanogaster* time point 1 to each other species time point 1, and then found the time points that minimized the distance between cell type profiles. The matching trajectory is shown by the blue line. The concordance between *D. melanogaster* and *D. simulans* morphological and expression profiles is high, while *D. yakuba* and *D. pseudoobscura* show a similar warping of time. Beyond time point 3, the matching between *D. melanogaster* and *D. virilis* becomes ambiguous, presumably reflecting the larger phylogenetic distance between these species. (B) By plotting the average all species-against-all species Euclidian distance between the cell type profiles, we show that the cell type profiles increase in similarity over developmental time, with a strong drop after time point 2. The thick line indicates the average distance between species, and the shaded area corresponds to the mean +/− the standard error of the mean. This result is consistent with the hourglass hypothesis of development, since the phylotypic period of *Drosophila* development is ~6 hours after time point 6.

The results of this analysis show that the time point (tp) matching closely follows the expected phylogenetic relationships between species. For *D. melanogaster* to *D. simulans*, the path closely follows the diagonal, though D. *melanogaster* tp5 matches *D. simulans* tp4 and *D. melanogaster* tp6 matches *D. simulans* tp5, indicating that *D. simulans*’ gene expression patterns are somewhat lagging behind the morphological progression. *D. yakuba* and *D. pseudoobscura* show similar patterns, in which *D. melanogaster* tp2 matches *D. yakuba/D. pseudoobscura* tp3, and *D. melanogaster* tp3-5 match *D. yakuba/D. pseudoobscura* tp4, indicating that the gene expression patterns in *D. yakuba* and *D. pseudoobscura* are progressing more slowly than the morphological marker in early time points. There is not a straightforward path to match *D. melanogaster* time points to *D. virilis* time points – particularly starting at time point 3, where there is not much difference in distance between *D. virilis* tp4-6, which mirrors the large phylogenetic distance between the two species. Therefore, there is not a strict matching between time points as determined by morphological markers and time points determined by cell type patterns.

Inspired by the hourglass model of development, we also hypothesized that variance in cell type fractions would decrease over time, as the phylotypic period of Drosophila development is generally thought to be ~6 hours after this stage in development (Kalinka et al., 2010). To test this hypothesis, for each time point, we calculated the Euclidian distances between the cell type vectors for all species against all other species, which results in a total of 10 comparisons per time point, i.e. *D. melanogaster* vs. the other four species, *D. simulans* vs. the three remaining species, etc. We found a large decrease in the average distance after time point 2, indicating that our data is consistent with an hourglass model of development with a phylotypic period that succeeds the time window under study (Figure 6B).

## Discussion

Here we report the generation of gene expression atlases for two species of *Drosophila*. Combined with existing data, we now have atlases with cellular-resolution measurements for key anterior-posterior patterning genes for five species of *Drosophilids* spanning roughly 40 million years of evolution. The comparison of these atlases reveals that the cell types, as defined by combinations of gene expression, are conserved between these species, but the proportions of theses cell types and their dynamics over the hour of blastoderm-stage development vary. We find that the average divergence of these cell type profiles decreases over the hour of developmental time measured here and that there is divergence between the dynamics of the cell type patterns and the progression of cellular membrane invagination.

We expect that these data sets, particularly combined with cross-species RNA-seq and ChIP-seq (Paris et al., 2013; Paris et al., 2015), will be a useful resource for the community interested in modeling developmental gene regulatory networks. The three previously-published expression atlases have been used to model the evolution of enhancer function (Wunderlich et al., 2012; Wunderlich et al., 2015), perform sensitivity analysis of domains of the patterning network (Bieler et al., 2011), develop detailed models of *eve* enhancer function (Ilsley et al., 2013; Staller et al., 2015b), and model spatially-varying transcription factor binding, when combined with ChIP-seq data (Kaplan et al., 2011). FlyEx, a one-dimensional data set that includes measurements of mRNA and protein expression in the embryo, has allowed for the development of detailed thermodynamic and dynamical models of gene expression (Jaeger et al., 2004; Poustelnikova et al., 2004; Janssens et al., 2006; Segal et al., 2008; Pisarev et al., 2009; He et al., 2010) that have revealed principles of enhancer function and canalization in the embryo.

Emerging techniques that allow for the detection of many more transcripts with spatial resolution will further augment the utility of the *Drosophila* embryo as a model for studying the evolution of developmental gene regulatory networks. For example, cyro-sliced RNA-seq allows for the measurement of the entire transcriptome in ~10 bins of cells along the anterior-posterior axis in both wild-type and mutant embryos (Combs and Eisen, 2013; Combs and Eisen, 2017). Single-cell RNA-seq also allows for the measurement of the whole transcriptome, albeit with lower signal to noise, in single cells. To map single cells back to their spatial location in the embryo, researchers rely on existing *in situ* data. A recent study applied single-cell RNA-seq in stage 6 *Drosophila* embryos and used the same *D. melanogaster* atlas analyzed here to spatially reconstruct the embryo from single cells (Karaiskos et al., 2017). In addition, improvements in the multiplexed *in situ* hybridization and sequencing approaches are enabling the measurement of hundreds or thousands of genes per cell (Lee et al., 2014; Choi et al., 2016; Eng et al., 2017) and will provide a useful way to measure a larger number of transcripts and to measure co-variation between gene expression patterns.

Deciphering how the *Drosophila* embryo is patterned has revealed fundamental insights about the molecules that control development (Nüsslein-Volhard and Wieschaus, 1980; Wieschaus, 2016), the architecture of gene regulatory networks (Lawrence, 1992; Jaeger, 2011; Jaeger et al., 2012), how GRNs are encoded in regulatory DNA (modENCODE et al., 2010; Wunderlich and DePace, 2011; Gregor et al., 2014; Vincent et al., 2016), and how GRNs and regulatory DNA evolve (Lynch and Roth, 2011). Here, we add an additional viewpoint on this flagship system, revealing the quantitative conservation of gene expression over 40 million years, and providing resources to the community for future studies of evolution of this conserved GRN.

## Methods

### Embryo collection and fixation

The sequenced stains of both *D. simulans* (Dsim\[w]501) and *D. virilis* (Dvir\b[1]; tb[1] gp-L2[1]; cd[1]; pe[1]) were used for these experiments. Embryos were collected on molasses plates in population cages at 23°C. D. simulans embryos were typically collected for 5 hours, and D. virilis embryos for 8 hours. After collection, embryos were de-chorionated in 50% bleach for 3 minutes and fixed in 10 mL heptane and 2.5 mL 10% methanol-free formaldehyde for 25 minutes while shaking. The formaldehyde was removed and 100% methanol added. A hard 1-minute shake removed the vitteline membrane, and the embryos were rinsed 3 times in 100% methanol and 2 times in 100% ethanol. Before staining, embryos were rehydrated in PBS with 0.2% Tween and 0.2% TritonX-100 (which we will call PBT-Tx). The embryos were post-fixed in 5% formaldehyde, 20 minutes for D. simulans and 25 minutes for D. virilis. They were then washed in PBT-Tx and then transferred to a hybridization buffer (5x SSC buffer, pH 4.2, 50% formamide, 40 μg/mL heparin, 100 μg/mL salmon sperm DNA, 0.2% TritonX-100) and incubated at 56°C for 1-6 hours.

### Probe synthesis and in situ hybridization

Species-specific RNA probes cloned into the pGEM-T Easy vector (Progema A1360) using the source cDNA or gDNA libraries and primers listed in Table S3. Probe synthesis was carried out as in (Fowlkes et al., 2011), using *in vitro* transcription of the probe template with either Sp6 or T7 RNA polymerase, depending on the orientation of the template in the pGEM-Teasy vector.

*In situ* hybridization reactions were carried as in (Fowlkes et al., 2011) with minor modifications. Briefly, ~100 μl of embryos were incubated for 24-48 hours at 56°C in 300 μl of hybridization buffer with 6 μl each of a DIG and DNP probe. We used a *ftz* DIG probe in each reaction as our fiduciary marker, and the DNP probe was for another gene of interest. Embryos were washed with stringent hybridization buffer (5x SSC buffer, 50% formamide, 0.2% TritonX-100) 10 times over 95 minutes at 56°C, and then blocked in 1% BSA in PBT-Tx for 1-2 hours. Probes were sequentially detected using horseradish-peroxidase (HRP) conjugated antibodies (anti-DIG POD, Sigma-Aldrich 11207733910 at 1:250; anti-DNP Perkin Elmer NEL747A001KT at 1:100) and either coumarin or Cy3 tyramide amplification reaction (Perkin-Elmer NEL703001KT, SAT704B). Between the DIG and DNP detection reactions, the anti-DIG HRP antibody was stripped by washing the embryos in stringent hybridization buffer 56°C and incubating them in 5% formaldehyde in PBT-Tx for 20 minutes. To remove all endogenous RNA, embryos were incubated in a 0.18 μg/ml RNAse A solution in PBT-Tx overnight at 37°C. Sytox Green (Life Technologies S7020, 1:5000) was used to stain the nuclei overnight at 4°C. To mount the embryos, embryos were dehydrated in solutions of increasing ethanol content and mounted in DePex (Electron Microscopy Service 13515) on a slide using 2 coverslips to create a bridge that prevents squashing of the embryos.

### Image acquisition and atlas generation

Z-stacks of embryos were acquired on a Zeiss LSM 710 with a plan-apochromat 20X 0.8NA objective at 1024×1024 pixels with 1 μm z-steps as described in (Fowlkes et al., 2011). Both RNA probe fluorophores (coumarin and Cy3) and the nuclear dye (Sytox Green) were excited at 750 nm, and the emitted light was split into three channels: 462–502 nm for coumarin, 514–543 nm for Sytox Green, and 599–676 nm. Using phase contract microscopy, embryos were staged using the percent membrane invagination as a morphological marker. Embryos were separated into 6 time points that correspond to 0-3%, 4-8%, 9-25%, 26-50%, 50-75% and 76-100% membrane invagination. The z-stacks were processed using previously-described software (Luengo Hendriks et al., 2006) which unmixes channels and segments individual nuclei to generate a pointcloud file for each embryo. Pointcloud files contain the 3D coordinates and fluorescence levels for each nucleus in the embryo.

To generate gene expression atlases, we used previously described methods (Fowlkes et al., 2008). For each species and time point, we generated morphological models that contain an average number of nuclei and 3D positions of nuclei that match the measured average egg length, embryo shape and nuclear density patterns. To find matching nuclei between time points, nuclear motion was constrained minimize distance and maximize smoothness. To enable fine registration of individual embryos to the template, the average expression pattern of our marker gene, *ftz*, was also included in the template for each species and time point. Each embryo pointcloud was coarsely aligned to the template using a rigid-body transformation and isotropic scaling and then finely aligned using non-rigid warping of the embryo to align marker gene boundaries with the template. To compute expression values for each nuclei and time point, we averaged measurements across those nuclei in individual pointclouds that were closest after spatial registration. To minimize expression variance, gains and offsets were estimated for expression measurements in each pointcloud prior to averaging. Additional details are available in (Fowlkes et al., 2008).

### Calculation of surface area and density

Surface area was computed as the sum of areas of the triangular mesh faces defined by the neighbor relation information between nuclei recorded in each embryo pointcloud (Luengo Hendriks et al., 2006). This represents the area of a surface passing through the centers of the nuclei (rather than, e.g. the surface area of the egg shell). Local density was computed by defining a disk of 15 μm radius on the surface around each nucleus, and dividing the number of nuclei in this disk by its area (Luengo Hendriks et al., 2006). These local densities were mapped onto the atlas cylindrical coordinate system, resampled to a regular grid, and averaged over each cohort.

### Cell type analysis

For this analysis, we used the *D. simulans* and *D. virilis* atlases described here, the “r2” version of the *D. melanogaster* atlas, and updated versions of the *D. pseudoobscura* and *D. yakuba* atlases, which now contain hb protein data (see Data Availability for details). We used a species-, and time point-specific threshold to distinguish nuclei that are “on” or “off” for each gene. The threshold is equal to the mode(values) + standard deviation(values), where values are the expression levels for a particular gene at one time point in one species. We have used this threshold calculation in several previous papers (Wunderlich et al., 2012; Staller et al., 2015a; Staller et al., 2015b) and finds that it effectively separates nuclei that are “on” from those with background levels of signal.

For the cell type analysis, we only considered the 9 genes measured in all species at all time points: *gt, hb [mRNA], kni, Kr, hkb, tll, eve, ftz*, and *odd*, using the thresholded data. In each nucleus at each time point, we identified the combination of genes expressed. To remove combinations that are uncommon and may be the result of measurement error, we eliminated combinations present in less than 0.1% of nuclei.

To calculate the similarity of the distributions of cell types between species and time points, we described each species time point as a 79×1 vector with each entry as the proportion of nuclei falling into each cell type. We calculated the Euclidian distances between these vectors. To match time points from *D. melanogaster* to each other species, we started by matching time point 1 in *D. melanogaster* to time point 1 in the other species, and for each subsequent *D. melanogaster* time point, we found the time point in the other species that had the minimum distance, while not allowing steps backwards in time. For example, if *D. melanogaster* time point 3 matched *D. yakuba* time point 4, *D. melanogaster* time point 4 can only match *D. yakuba* time points 4, 5 or 6.

### Clustering analysis

For the clustering analysis, we described each species/time point combination as a 79×1 vector with each entry as the proportion of nuclei corresponding to each cell type. We then normalized each row (cell type) by the mean. We performed hierarchical clustering of the mean-centered data using Euclidian distance as the distance measure and average linkage.

### Calculation of expression distance score

Expression levels for each gene were scaled so that the maximum expression level across all cells was 1 at each time point. Cell-to-cell expression profile comparisons between species atlases were computed using the squared Euclidian distance between the vectors of average expression measurements for the cell across all 6 time points and the 9 genes measured in all species at all time points *(hb* [protein*], gt, kni, Kr*, *tll, hkb*, *eve, odd* and *ftz)*. We also performed the matching using *hb* [mRNA], but found the hb [protein] data gave more closely matching cells, presumably because the mRNA expression pattern is less conserved than the protein expression pattern.

For a pair of nuclei *i* and *j* this distance was given by:

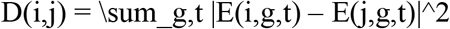

where ***E***(***i,g,t***) is the expression of the ***g****th* gene recorded in the atlas for the ***i**th* cell at time point ***t***. We utilized squared distance since it is additive across genes and time-points, making the contribution of individual genes more interpretable. Comparisons were only made to cells in corresponding regions of the embryo. Corresponding locations were estimated by scaling each atlas to unit egg length and nearby nuclei were specified as those nuclei in the target embryo that were within the 30 nearest to the cell to be matched. To visualize displacement to the best match, we used the average of the 3D locations of the 10 cells with the smallest expression distance, weighted by the inverse of their expression distance. This provides a more stable estimate of displacement when there are multiple good matching cells. The 3D displacement estimates were visualized in cylindrical projection.

## Data Availability

The *D. simulans* (release 1, r1) and *D. virilis* (r1) atlases are available on FigShare (10.6084/m9.figshare.6866795). Also available on FigShare are updated *D. yakuba* (r2) and *D. pseudoobscura* (r2) atlases, which now contain hb protein expression data. The *D. melanogaster atlas* is unchanged from the version (r2) available here: http://bdtnp.lbl.gov/Fly-Net/bidatlas.jsp.

## Acknowledgements

We thank John Reinitz for providing the hunchback antibody, the UC San Diego (now Cornell) Drosophila Stock Center for the *D. simulans* and *D. virilis* stocks, and Cris Luengo and Lisa Simirenko for assistance with the image processing pipeline. We also thank Peter Combs for helpful comments on the manuscript.

---

Figure S1. This figure shows the distributions of expression distance scores for Figure 5, in the case of matching nuclei to their nearest neighbor (left column) or to their best local neighbor (right column). In each species, compared to *D. melanogaster*, the local search appreciably decreases the mean and median expression distance score, with the most largest decreases in the *D. melanogaster-D. pseudoobscura* and *D. melanogaster-D. virilis* comparisons.

**Table S1.** Number of embryos per cohort in *D. simulans* and *D. virilis* datasets

| Gene | Species | 5:0-3% | 5:4-8% | 5:9-25% | 5:26-50% | 5:52-75% | 5:76-100% |
| --- | --- | --- | --- | --- | --- | --- | --- |
| bcd | D. sim | 9 | 6 | 1 |  |  |  |
| cad | D. sim | 13 | 6 | 11 | 5 | 4 | 6 |
| hb | D. sim | 5 | 5 | 3 | 3 | 4 | 3 |
| hb [protein] | D. sim | 9 | 16 | 15 | 4 | 1 | 12 |
| gt | D. sim | 5 | 7 | 6 | 7 | 6 | 5 |
| Kr | D. sim | 9 | 10 | 10 | 7 | 12 | 13 |
| kni | D. sim | 7 | 7 | 15 | 8 | 5 | 5 |
| fkh | D. sim | 5 | 11 | 13 | 12 | 5 | 6 |
| hkb | D. sim | 3 | 9 | 8 | 10 | 7 | 7 |
| tll | D. sim | 5 | 7 | 9 | 7 | 5 | 7 |
| eve | D. sim | 6 | 11 | 12 | 8 | 4 | 10 |
| ftz | D. sim | 9 | 11 | 19 | 11 | 7 | 17 |
| odd | D. sim | 11 | 14 | 13 | 9 | 4 | 9 |
| prd | D. sim | 12 | 4 | 14 | 12 | 6 | 8 |
| bcd | D. vir | 4 | 5 |  |  |  |  |
| hb | D. vir | 12 | 12 | 7 | 5 | 6 | 13 |
| hb [protein] | D. vir | 3 | 4 | 6 | 4 | 2 | 8 |
| gt | D. vir | 4 | 5 | 7 | 1 | 4 | 4 |
| Kr | D. vir | 5 | 11 | 7 | 6 | 4 | 6 |
| kni | D. vir | 13 | 12 | 8 | 4 | 2 | 15 |
| hkb | D. vir | 5 | 4 | 2 | 3 | 4 | 4 |
| tll | D. vir | 9 | 10 | 7 | 3 | 4 | 5 |
| eve | D. vir | 10 | 13 | 10 | 7 | 9 | 22 |
| ftz | D. vir | 4 | 11 | 9 | 9 | 9 | 23 |
| odd | D. vir | 6 | 8 | 7 | 6 | 7 | 18 |
| hb [protein] | D. pse | 13 | 5 | 4 | 2 | 1 | 7 |
| hb [protein] | D. yak | 11 | 12 | 8 | 10 | 4 | 9 |

**Table S2.**
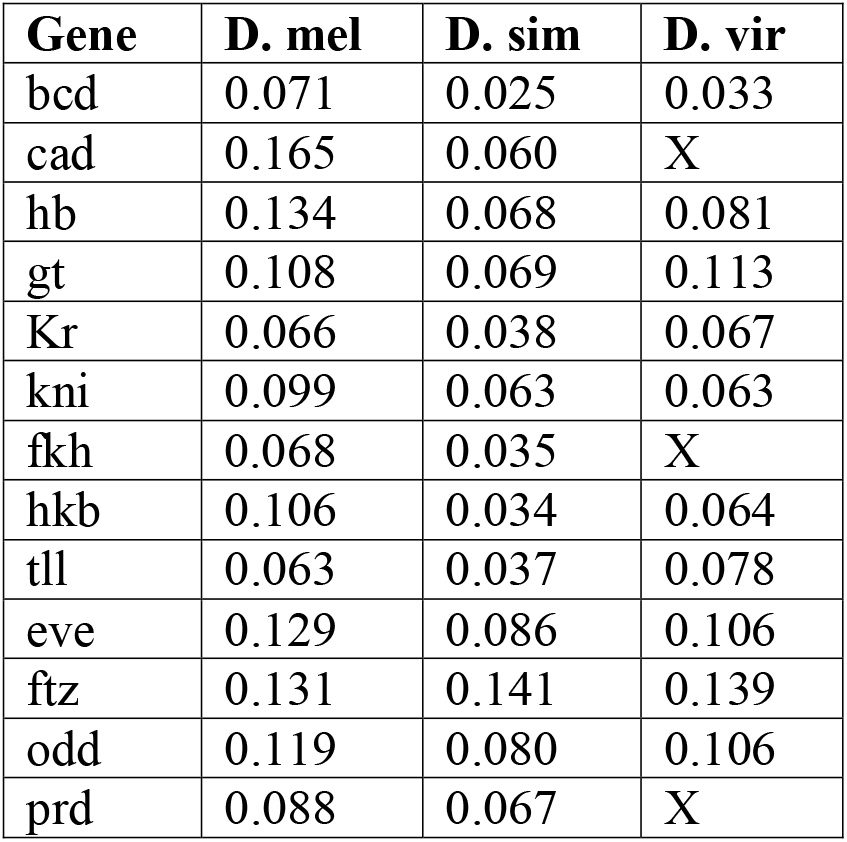
Average standard deviation across cohorts

**Table S3.**
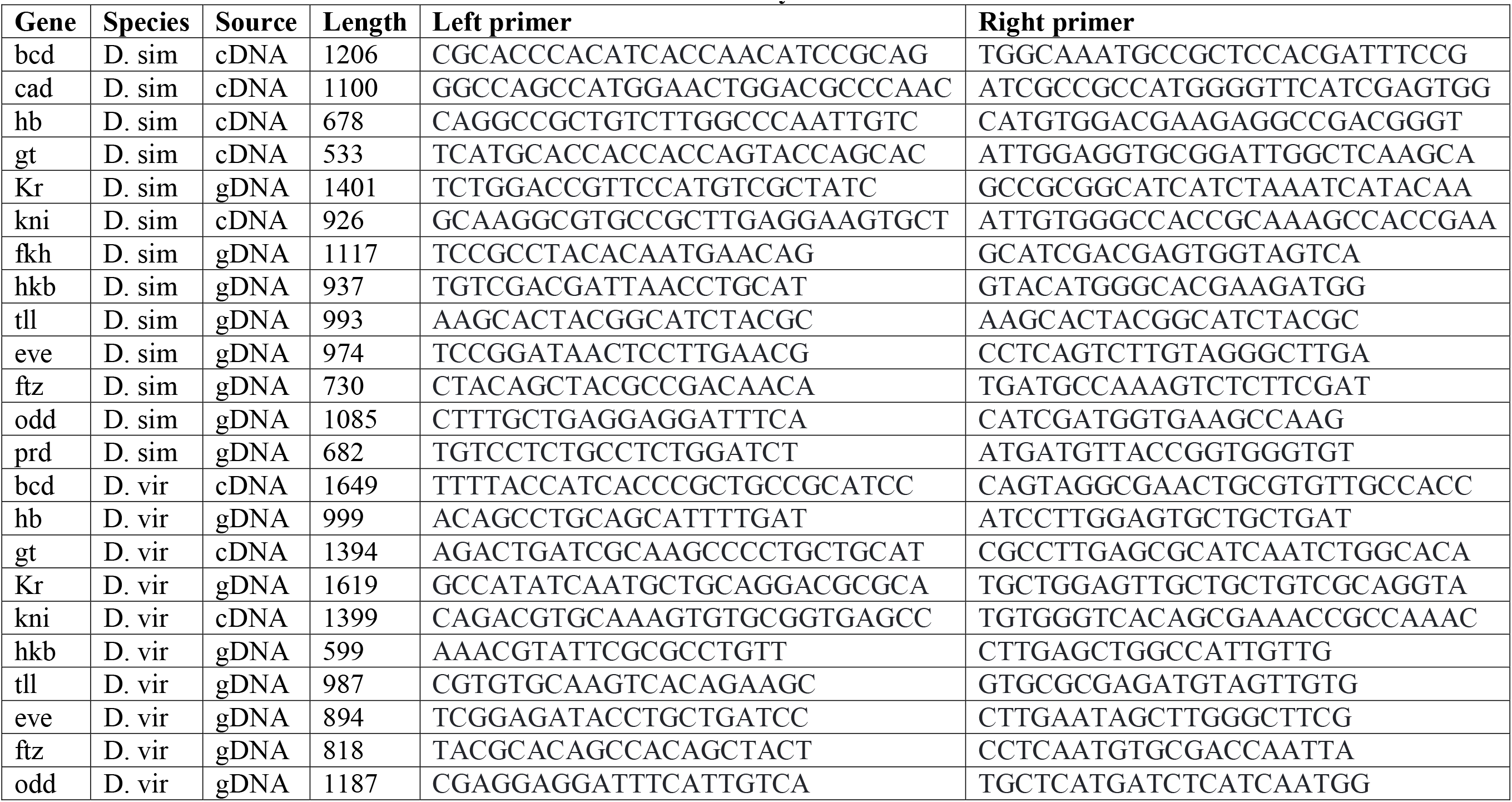
Probe constructs for *D. simulans* and *D. virilis in situ* hybridizations

